# Short hairpin RNAs artifactually impair cell growth and suppress clustered microRNA expression

**DOI:** 10.1101/372920

**Authors:** JT Powers, EL da Rocha, DS Pearson, P Missios, TY de Soysa, J Barragan, P Cahan, GQ Daley

## Abstract

Functional gene disruption is a central tenet of cancer research, where novel drug targets are often identified and validated through cell-growth based knockdown studies or screens. Short hairpin RNA (shRNA)-mediated mRNA knockdown is widely used in both academic and pharmaceutical settings. However, off-target effects of shRNAs as well as interference with endogenous small RNA processing have been reported. We show here that lentiviral delivery of both gene-specific and non-targeting control shRNAs impair *in vitro* cell growth in a sequence independent manner. In addition, exogenous shRNAs induce a depressed cell-cycle-gene expression signature that is also shRNA-sequence independent and present across several studies. Further, we observe an shRNA mediated general repression of microRNAs belonging to polycistronic genetic clusters, including microRNAs from established oncogenic microRNA clusters. The collective impact of these observations is particularly relevant for cancer research, given the widespread historical use of shRNAs and the common goal of interrogating genes that regulate proliferation. We therefore recommend that when employing shRNA for target validation, care be taken to titrate shRNA dose, use hairpin-expressing controls, perform gene-of-interest rescue experiments and/or corroborate shRNA-derived results by small interfering RNA (siRNA) knockdown or CRISPR/Cas9-mediated genetic knockout. Minimizing these deleterious sequence independent effects will improve research fidelity and help address reported challenges in experimental reproducibility.

---

Functional gene disruption is a central tenet of cancer research, where novel drug targets are often identified and validated through cell-growth based knockdown studies or screens. Short hairpin RNA (shRNA)-mediated mRNA knockdown is widely used in both academic and pharmaceutical settings. However, off-target effects of shRNAs as well as interference with endogenous small RNA processing have been reported^1–3^. We show here that lentiviral delivery of both gene-specific and non-targeting control shRNAs impair *in vitro* cell growth in a sequence independent manner. In addition, exogenous shRNAs induce a depressed cell-cycle-gene expression signature that is also shRNA-sequence independent and present across several studies. Further, we observe an shRNA mediated general repression of microRNAs belonging to polycistronic genetic clusters, including microRNAs from established oncogenic microRNA clusters. The collective impact of these observations is particularly relevant for cancer research, given the widespread historical use of shRNAs and the common goal of interrogating genes that regulate proliferation. We therefore recommend that when employing shRNA for target validation, care be taken to titrate shRNA dose, use hairpin-expressing controls, perform gene-of-interest rescue experiments and/or corroborate shRNA-derived results by small interfering RNA (siRNA) knockdown or CRISPR/Cas9-mediated genetic knockout. Minimizing these deleterious sequence independent effects will improve research fidelity and help address reported challenges in experimental reproducibility^4^.

Gene-targeting shRNAs are typically introduced into cell lines *in vitro* through lentiviral infection, leading to genomic integration and permanent expression of the hairpin construct. Functionally similar to siRNAs, which are directly loaded into the RISC complex, shRNA must first be transcribed and processed by cellular microRNA biogenesis machinery. ShRNAs are valued because they are easy to use and can affect long term knockdown of a gene, whereas siRNA knockdown is transient. CRISPR/Cas9-mediated knockout also results in a permanent genetic event, though in some cases knockdown studies are preferred to evaluate intermediate gene expression phenotypes and avoid severe phenotypes or lethality due to complete loss of gene function. While shRNA knockdown occupies an important niche between short term and permanent gene disruption, their use comes with caveats. In addition to off-target effects and impact on small RNA processing machinery, shRNA-induced liver and neuronal toxicity have been documented *in vivo*^3,5^. In addition, shRNA-mediated cytotoxicity has been implicated in *MYC*-driven hepatocellular carcinoma in mice^6^.

In our recent work on the role of *LIN28B* and its microRNA target *let-7* in neuroblastoma we described a reduced-growth phenotype produced by shRNA-mediated *LIN28B* knockdown that was not replicated by either siRNA or CRISPR/Cas9-mediated *LIN28B* knockout^7^. To further explore this observed shRNA induced toxicity, we first compared the effects of shRNA-and siRNA-mediated *LIN28B* knockdown on the cell cycle profile of BE(2)C neuroblastoma cells (*Fig. 1a, b*). We observed disturbed profiles in both non-targeting control and *LIN28B*-targeting shRNA infected cells (*Fig. 1c*). This effect was not observed in control or *LIN28B*-siRNA transfected cells, consistent with our previous work (*Fig. 1d*). *MYCN*-siRNA served a control for an altered cell cycle profile (*Fig. 1d*). We next tested this shRNA/siRNA relationship in K562 cells, a morphologically distinct leukemia cell line that also expresses *LIN28B*, and again observed that multiple *LIN28B*-targeting shRNAs reduced cell growth similar to the *ABL1* siRNA control, while two *LIN28B*-targeting siRNAs did not (*Fig. 1e, f*). We observed a similar pattern of shRNA, but not siRNA, induced growth impairment during the course of studies of the *NAT10* acetyltransferase in BE(2)C cells (*Supp. Fig. 1a, b*). *LIN28B* has recently been implicated in pancreatic cancer, where pancreatic cell growth inhibition was demonstrated using *LIN28B*-specific shRNAs^8^. However, we observed that the same pancreatic cancer cell lines were entirely refractory to complete LIN28B protein loss mediated by CRISPR/Cas9 targeting of the *LIN28B* genetic locus with multiple sgRNAs (*Supp. Fig. 2a, b*). Together, these results suggest that exogenous shRNA expression may generally suppress cell growth independent of hairpin sequence, potentially leading to misleading experimental conclusions. Of note is the observation that the non-targeting control hairpin induced the same cell cycle defect as gene-targeting shRNAs (*Fig. 1c*).

**Figure 1.**
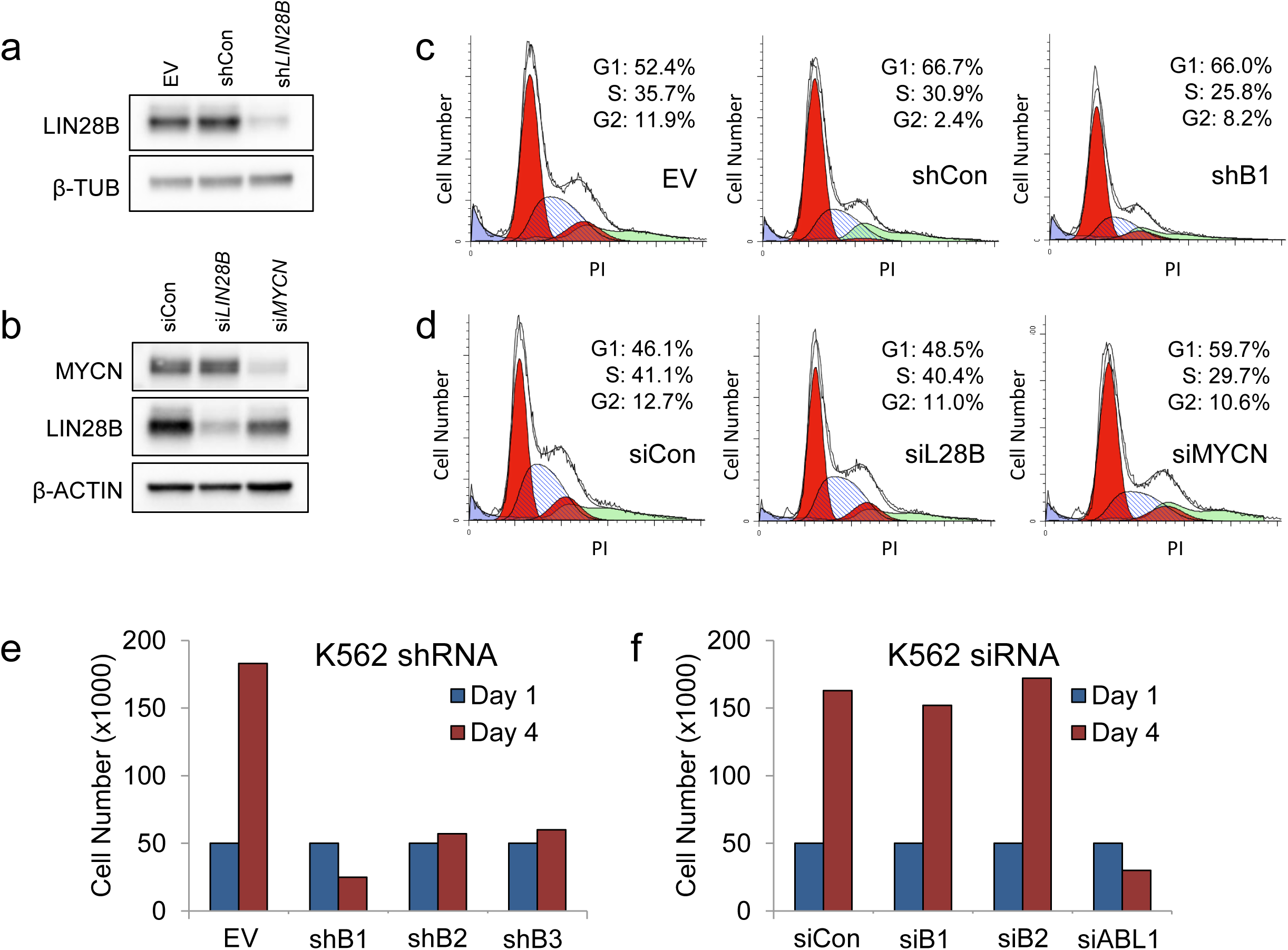
shRNA expression impairs cell growth. **(a,b)** Immunoblot for LIN28B in BE(2)C neuroblastoma cells transduced with indicated lentivirus (a) or siRNA (b) for 2 days. **(c,d)** Cell cycle profiles of PI stained cells treated as in *a* and *b*, respectively. **(e)** Growth analysis of K562 leukemia cells infected with the indicated lentivirus (MOI 20). **(f)** Cell growth analysis of K562 cells transfected with the indicated siRNA. (EV, empty vector; Con, non-targeting shRNA control; B1/B2/B3, *LIN28B* targeting shRNAs)

**Figure 2.**
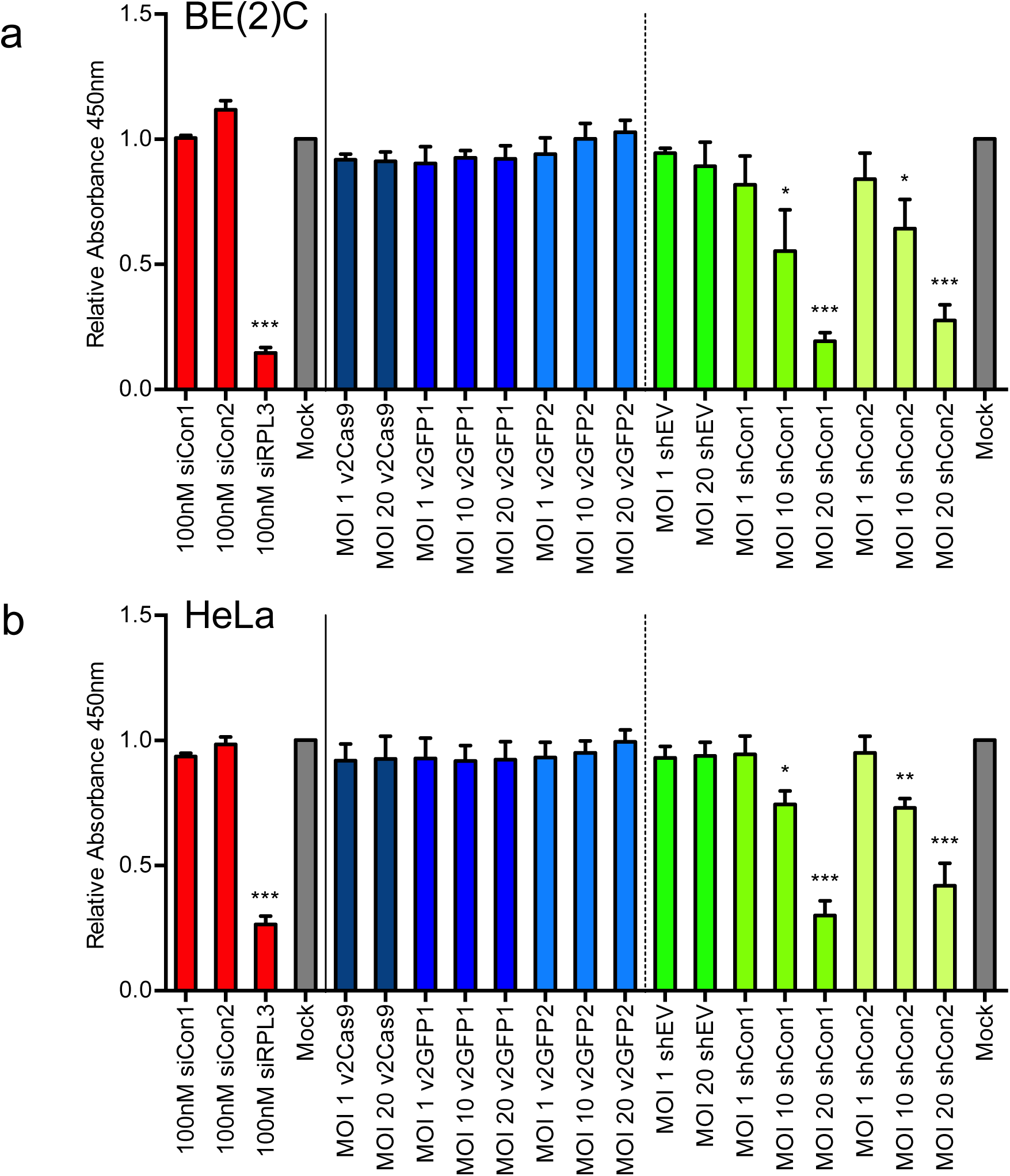
Control shRNAs inhibit cell growth. **(a,b)** Relative cell growth in BE(2)C and HeLa cells five days after the indicated transfections or infections. Dose for each treatment is noted. Values are relative to sham transfections or infections, respectively. (*p<0.05, **p<0.01, ***p<0.001, unpaired t-test).

To further understand the apparent sequence independent growth inhibition of shRNAs, we interrogated the effect of a panel of scrambled siRNA, *GFP*-targeting CRISPR/Cas9, and non-targeting shRNA controls on the growth of BE(2)C and HeLa cells five days post treatment. This approach allowed us to test each platform for effects on proliferation that were independent of unintentional gene targeting side effects. We intentionally used a high concentration of siRNA (100nM), as well as an siRNA against *RPL3,* a target well documented to impair cell proliferation^9^ as a positive control for reduced cell growth. Multiplicity of infection (MOI) for lentiviral CRISPR/Cas9 and shRNA constructs ranged from low MOIs of 1 to high MOIs of 20. Neither scrambled sequence control siRNAs displayed growth abnormalities, similar to the mock-transfection, whereas the *RPL3*-targeting siRNA severely impaired cell growth in both cell types, as expected (*Fig. 2a,b*). Cas9 and Cas9/*GFP*-targeting-sgRNA infected cells responded similarly, with neither construct perturbing growth compared to mock-infection, regardless of MOI. In contrast, scrambled sequence shRNA-expressing lentiviruses significantly reduced both BE(2)C and HeLa cell proliferation over the 5 day experiment, in a dose-dependent manner at MOIs of 10 and 20 (*Fig. 2a, b*).

We sought to gain insight into the observed sequence non-specific shRNA induced proliferative defect and performed global mRNA expression analysis on BE(2)C cells infected with empty lentivirus, non-targeting control shRNA, or each of three *LIN28B* targeting shRNAs. Replicate samples showed high correlation within each condition (*Fig. 3a*). Given the proliferation phenotype driven by shRNAs, we first identified the cell cycle genes that were differentially regulated after shRNA transduction and noted that all four shRNA samples, again including the non-targeting control, displayed significant down regulation of cell cycle genes compared to empty vector (*Fig. 3b*). Furthermore, the most significantly effected Gene Ontology processes were almost universally cell cycle related in all four shRNA samples (*Fig. 3c*). These data are consistent with a model wherein high level exogenous shRNA expression has a sequence-independent effect on cell proliferation as demonstrated by consistent growth impairment and downregulation of cell cycle related processes (*Fig. 2, Fig. 3c*).

**Figure 3.**
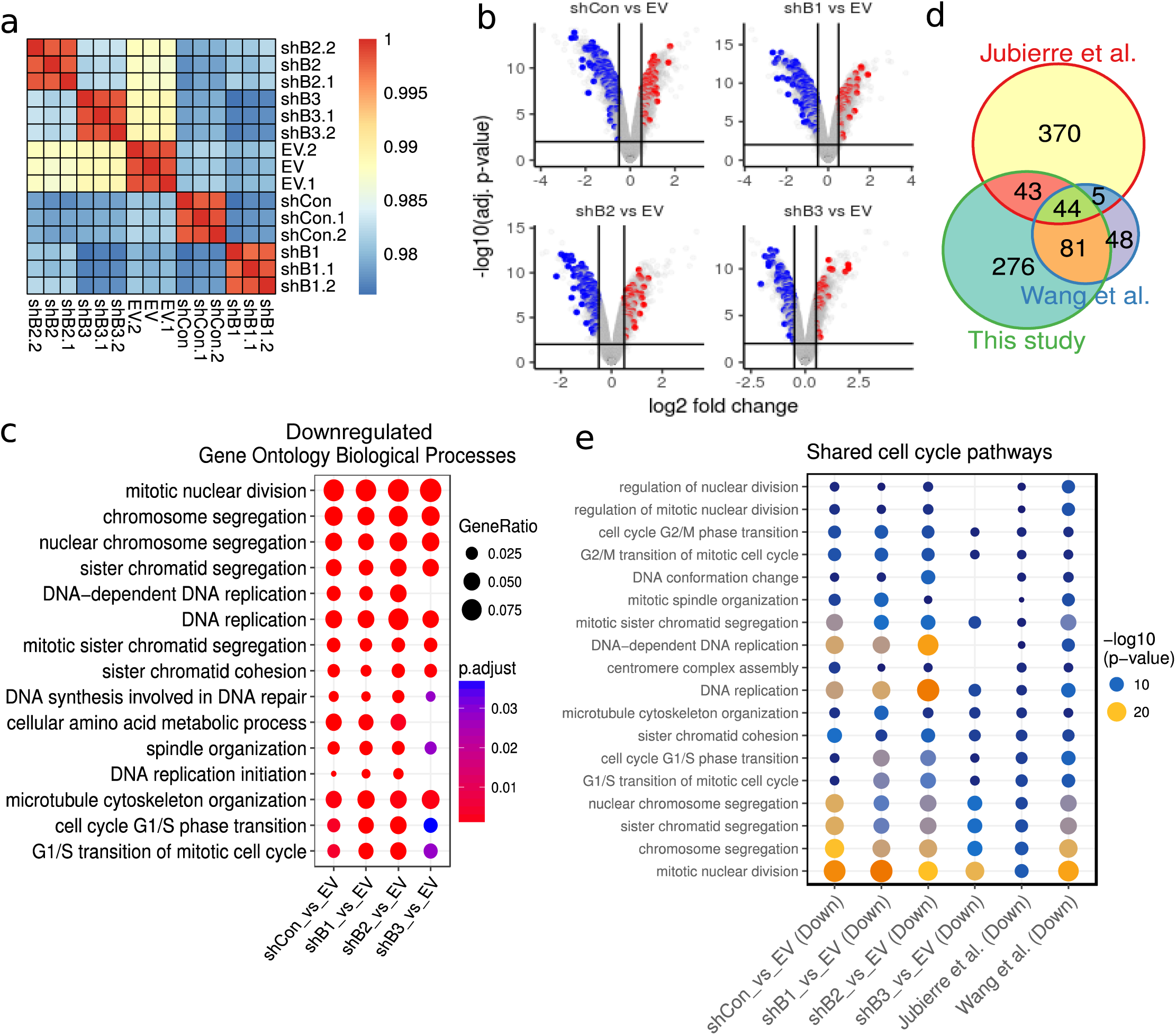
Effect of exogenous shRNAs on gene expression patterning. **(a)** Clustering of Pearson correlation coefficients between gene expression profiles of BE(2)C cells infected with the indicted lentiviral constructs. **(b)** Volcano plots colored by differentially expressed cell cycle-related genes. Blue indicates downregulation; Red indicates upregulation. **(c)** Gene Ontology analysis of genes significantly downregulated in each comparison. **(d)** Venn diagram of downregulated gene ontology biological processes between our dataset and datasets generated by Jubierre *et al.* and Wang *et. al.* **(e)** Gene ontology biological processes downregulated across the studies in *d*.

We compared our expression data to two existing shRNA-based microarray datasets; one from BE(2)C neuroblastoma cells and the other from DU145 prostate cancer cells^10,11^. Correlative analysis within the two studies showed agreement within sample replicates (*Supp. Fig. 3a, 4a*). Differential gene expression analysis showed a preferential down-regulation of cell cycle genes, consistent with our own data (*Supp. Fig. 3b, 4b*). Gene Ontology analysis of the most significantly altered pathways also revealed a strong cell cycle signature in both datasets (*Supp Fig, 3c, 4c).* Indeed, Venn diagram overlap of downregulated Gene Ontology processes revealed 44 common to all the three datasets (*Fig. 3d; Supp. Table 1*). Of note, 42 of them were cell cycle processes. Moreover, the 18 most highly significant for each comparison were all cell cycle-related and showed strong agreement across the studies. Note the agreement between our control-shRNA pattern and those of the other two studies (*Fig. 3e*). These inter-study observations are consistent with our own data and support a model where small hairpin RNAs inhibit cell growth in a sequence-independent manner.

Following transcription, shRNAs are processed by the microRNA biogenesis machinery and have been reported to impact microRNA levels^3,12,13^. We therefore examined global microRNA expression in the same set of BE(2)C shRNA transduced cells. Analysis of microRNA expression patterns revealed high correlation among all of the shRNA samples, including the non-targeting scrambled sequence control shRNA, which was more closely correlated with *LIN28B* shRNA samples than empty vector control. This pattern is consistent with observed sequence-independent effects on proliferation (*Fig. 1, Supp. Fig. 1, Fig.4a*). Of the 50 most downregulated microRNAs in each sample set, 25 were consistent between control shRNA and all three *LIN28B* shRNA-expressing cells (*Fig. 4b*). Upon analysis of these 25 microRNAs, we discovered that 24 of them belonged to clustered microRNA loci (*Fig. 4c*). Moreover, when we examined expression of clustered *vs*. single-microRNA loci across the genome, we observed a strong global downregulation of clustered microRNA expression compared to singleton microRNAs (*Fig. 4d*). These data indicated that the broad cell growth impairment observed following transduction of high levels of shRNA correlated markedly with aberrant processing of clustered species of microRNA.

**Figure 4.**
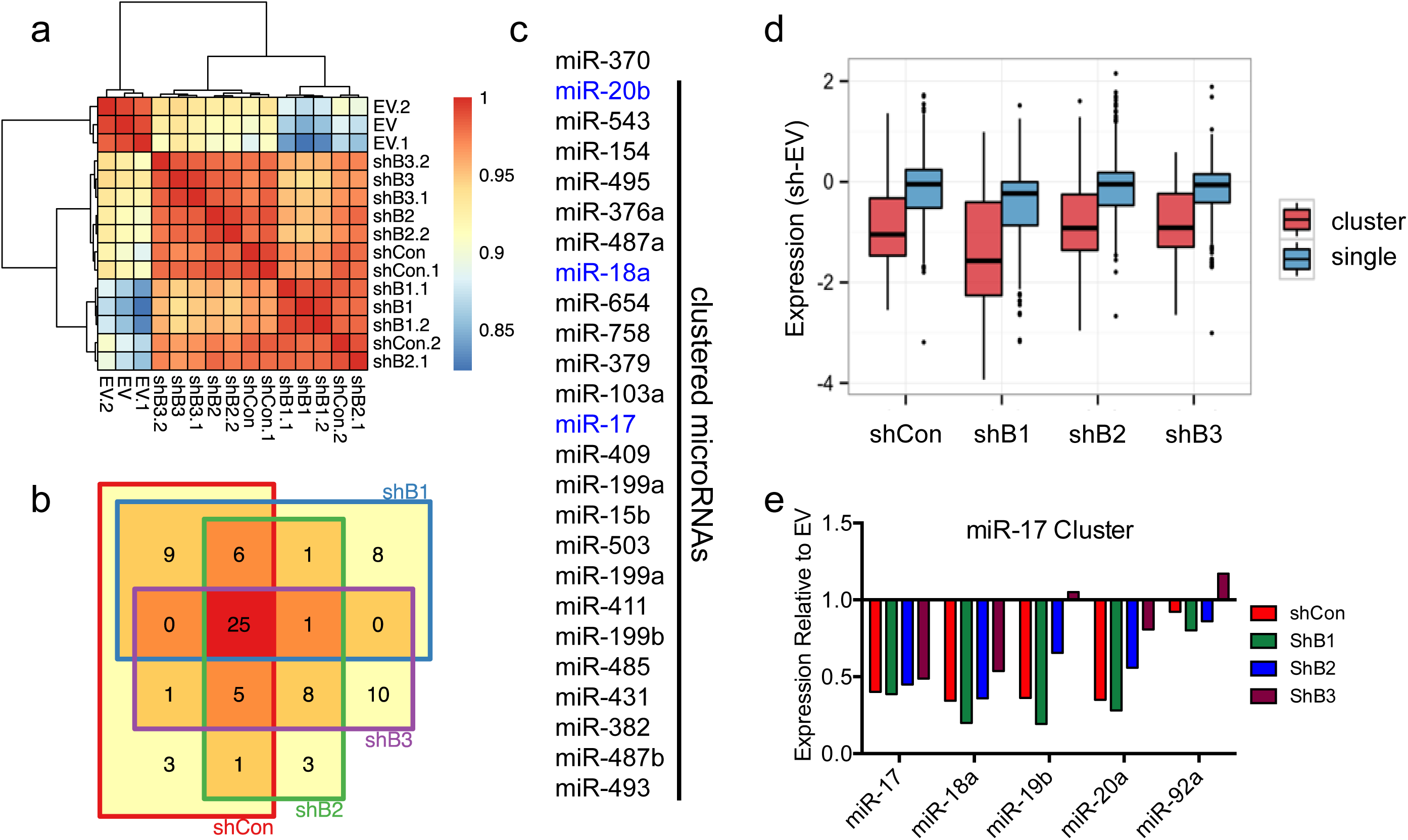
Effect of exogenous shRNAs on microRNA expression patterning. **(a)** Clustering of Pearson correlation coefficients between gene expression profiles of BE(2)C cells infected with the indicated shRNA constructs. **(b)** Venn diagram of the 50 most downregulated microRNAs in each the BE(2)C infections from *a.* **(c)** List of the 25 commonly downregulated microRNAs from *b*. OncomiRs are in blue. **(d)** Relative microRNA expression from clustered or singleton loci in cells from *a.* Values are relative to EV. **(e)** Analysis of relative miR-17 OncomiR cluster expression in cells from *a*. Values are relative to empty vector control. (EV, empty vector; Con, non-targeting shRNA; B1/B2/B3, *LIN28B* targeting shRNAs)

The microRNAs of the *miR-17, miR-18, miR-19*, and *miR-25* families are highly expressed in multiple cancers. They are established pro-growth oncogenes, and are all transcribed from one of three OncomiR clusters^14,15^. Indeed, eight of these microRNAs are among the highest expressed microRNAs in BE(2)C cells (*Supp. Fig. 5a*). The *miR-17-92* cluster is the best characterized of these and has been identified as a transcriptional target of *MYC*^16^. Upon shRNA infection of BE(2)C cells, microRNA levels from the *miR-17-92* OncomiR cluster were broadly suppressed in all shRNA samples, including non-targeting control shRNA (*Fig. 4e*). Further, we observed the same pattern of repression in the remaining two OncomiR clusters (*Supp Fig. 5c, d*). Given their importance in cancer biology, sequence-independent shRNA-induced suppression of these oncogenic microRNA clusters may offer a plausible explanation for the proliferative defects observed in our study.

Loss of gene function analysis is indispensable to cancer research. Within the *Nature* family of journals alone, chosen for analysis due to their high rate of depositing published manuscripts into pubmed central, there have been over three thousand publications in cancer research over the last ten years, well over a thousand of which have included gene knock-down data (*Supp. Fig. 6a-c*). Tool fidelity in such a widely used research approach is therefore critical. We demonstrate here that shRNA can impact cell growth *in vitro* as well as preferentially disrupt clustered microRNA expression in a sequence-independent manner. We further identify a depressed cell cycle Gene Ontology signature that is consistent across multiple shRNA studies. Our data poses a significant concern for use of shRNA knockdown for cancer research in particular, where the common goal is to identify genes whose knock-down compromises cell growth. This is particularly true for easy-to-infect cells, which include most adherent lines. Difficult-to-infect cells, such as lymphocytes, might be inherently amenable to shRNA use, due to their self-limitation of viral dose^17^. Indeed, Sigma-Aldrich manufacturer protocol suggests using and MOI of 1 to deliver shRNA virus to cells. When we infected BE(2)C cells with *LIN28B* shRNA virus below an MOI of 1, two out of three failed to efficiently knockdown LIN28B expression (*Supp. Fig. 7*). The challenge therefore is to titrate viral dose only as high as needed to achieve target suppression while remaining below cytotoxic levels.

To address the shortcomings of small hairpin RNAs and to reduce potential contributions to irreproducibility in cancer research, we recommend the following practices for shRNA use. Care should be taken to titer viral supernatants, followed by use of the lowest MOI possible to effectively knockdown the gene of interest. Non-targeting scrambled sequence hairpin controls should always be used at similar MOI to control for the sequence-independent effects reported in this manuscript. An additional empty vector control should also be used to control for general effects of shRNA expression. Finally, phenotypes observed through effective use of shRNA knockdown should be reverted by expression of shRNA-resistant versions of the gene of interest and/or validated by either siRNA or CRISPR/Cas9 use (*Supp. Fig. 6d, 8*).

Reproducibility is central to scientific advancement, but has been difficult to achieve in recent years, especially in the field of cancer research^4,18^. Research tools such as exogenous shRNA that inappropriately effect the same cell growth inhibition that is frequently the goal of target validation studies in cancer research have almost certainly contributed, despite good faith efforts of the researchers involved. Minimizing shRNA-induced, sequence independent detrimental effects on cell growth and expression signatures will reduce false positive results and contribute to improved reproducibility in cancer and potentially other research fields.

## Materials and Methods

### Cell Culture

BE(2)C (ATCC CRL-2268), HeLa (ATCC CCL-2), and K562 (ATC CCL-243) were maintained in RPMI media with 10% heat-inactivated fetal calf serum, 1 μg ml-1 penicillin, and 1 U ml-1 streptomycin. All cell lines were purchased for the purposes of this study, are not among commonly misidentified cell lines (according to the International Cell Line Authentication Committee), and tested negative for mycoplasma contamination.

### siRNA

siRNA transfections were performed with Lipofectamine™ RNAiMAX Transfection Reagent as per the manufacturer protocol. siRNAs used: siCon/Con1 (ThermoFisher cat# 4390843); siCon2 (ThermoFisher cat# 4390846); si*LIN28B* (ThermoFisher cat# 4392420, identifier s52477); si*MYCN* (ThermoFisher cat# 4392420, identifier s9134); si*ABL1* (ThermoFisher cat# 4390824, identifier s864); si*RPL3* (ThermoFisher cat# 4392420, identifier s12142); si*NAT10*-1 (ThermoFisher cat# 4392420, identifier s30491); si*NAT10*-2 (ThermoFisher cat# 4392420, identifier s30492).

### Lentivirus

Lentiviral shRNA particles were prepared as previously described^19,20^. Viral titers were determined using Takara Lenti-X™ GoStix™ as per the manufacturer protocol. *ShRNA constructs used*: shCon/Con1 (SHC216, sequence 5’-CGTGATCTTCACCGACAAGAT-3’); shCon2 (SHC204, sequence 5’-CGTGATCTTCACCGACAAGAT-3’); shEV (SHC201-no insert); shLIN28B1/B1 (TRCN0000144508, sequence 5’-CCTGTTTAGGAAGTGAAAGAA-3’); shLIN28B2/B2 (TRCN0000122599, sequence 5’-GCCTTGAGTCAATACGGGTAA-3’); shLIN28B3/B3 (TRCN0000122191, sequence 5’-GCAGGCATAATAAGCAAGTTA-3’); shNAT10-1 (TRCN0000296411, sequence 5’-TTGCTGTTCACCCAGATTATC-3’); shNAT10-2 (TRCN0000035702, sequence 5’-CGCAAAGTTGTGAAGCTATTT-3’). *CRISPR/Cas9 constructs used:* V2-Cas9 (lentiCRISPRv2, Addgene plasmid #52961), V2-GFP1 was created by cloning eGFP sgRNA sequence 5’-GGGCGAGGAGCTGTTCACCG-3’ into the lentiCRISPRv2 vector. V2-GFP2 was created by cloning eGFP sgRNA sequence 5’-GAGCTGGACGGCGACGTAAA-3’ into the lentiCRISPRv2 vector. LentiCRISPRv2 was a gift from F. Zhang^19,20^.

### Western Blotting

Western blots were performed with antibodies against LIN28B (Cell Signaling 4196), MYCN (Santa Cruz Biotechnology sc-53993), NAT10 (Proteintech 13365-1-AP), β-Tubulin (Cell Signaling 2146), and β-Actin (Santa Cruz Biotechnology sc-8342).

### Raw data processing of microarray data

The Affymetrix GeneChip miRNA 3.0 arrays were processed in R as follows. First, we re-annotaed the probes we loaded raw.CEL files using the ‘oligo’ and ‘affy’ packages^21,22^. Second, we empirically defined a background level of expression as 5.41 by determining two standard deviations above the mean probe intensity of all probes mapping to Zea mays. We excluded all non-human probes and probes mapping to un-annotated genes, leaving 1,724 probes for further analysis. We further restricted downstream analysis to the 542 probes were detected (i.e. mean expression in the group > the background defined above) in at least one experimental group. We used the mirFocus database (http://mirfocus.org/index.php) to annotate miRNAs as clustered or singletons.

### Reanalysis of publicly available datasets

We reanalyzed publicly available microarray datasets. The accession numbers for theses studies are GSE46708 and GSE74622. For GSE46708, we downloaded processed data from GEO. For GSE74622, we used the R packages oligo and the annotation package pd.hugene.2.0.st to process raw.CEL files. Differential expression analysis was performed using the R package limma. We used the R packages clusterProfiler and ReactomePA for gene ontology enrichment analysis. For GSE46708, we used genes with log2 fold changes > 0.5 or < −0.5 for gene ontology analysis. For GSE74622, we used genes with log2 fold changes > 0.25 or < −0.25 for gene ontology analysis. Clustering and heat map visualization were performed the pheatmap package.

### Literature analysis

We downloaded xml files from ftp://ftp.ncbi.nlm.nih.gov/pub/pmc and wrote customized scripts to parse these xml files and query combinations of keywords throughout each manuscript. These scripts are available from the authors upon reasonable request.

## Figure Legends

**Figure S1.**
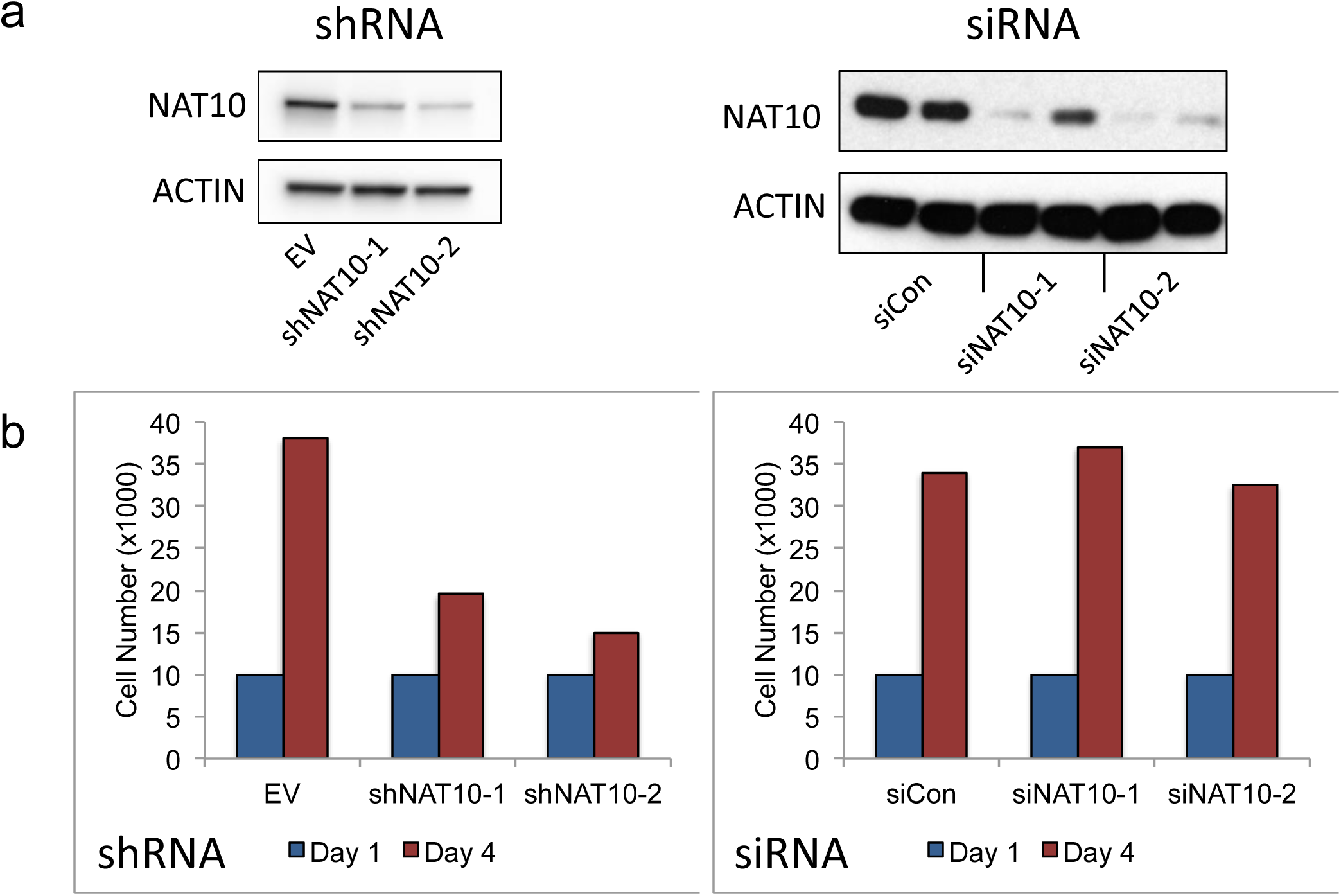
*NAT10* targeting shRNAs impair cell growth. **(a)** Immunoblot for NAT10 in BE(2)C cells infected or transfected with the indicated lentiviral constructs or siRNAs, respectively. **(b)** Cell growth analysis of BE(2)C cells treated as in *a*. (EV, empty vector)

**Figure S2.**
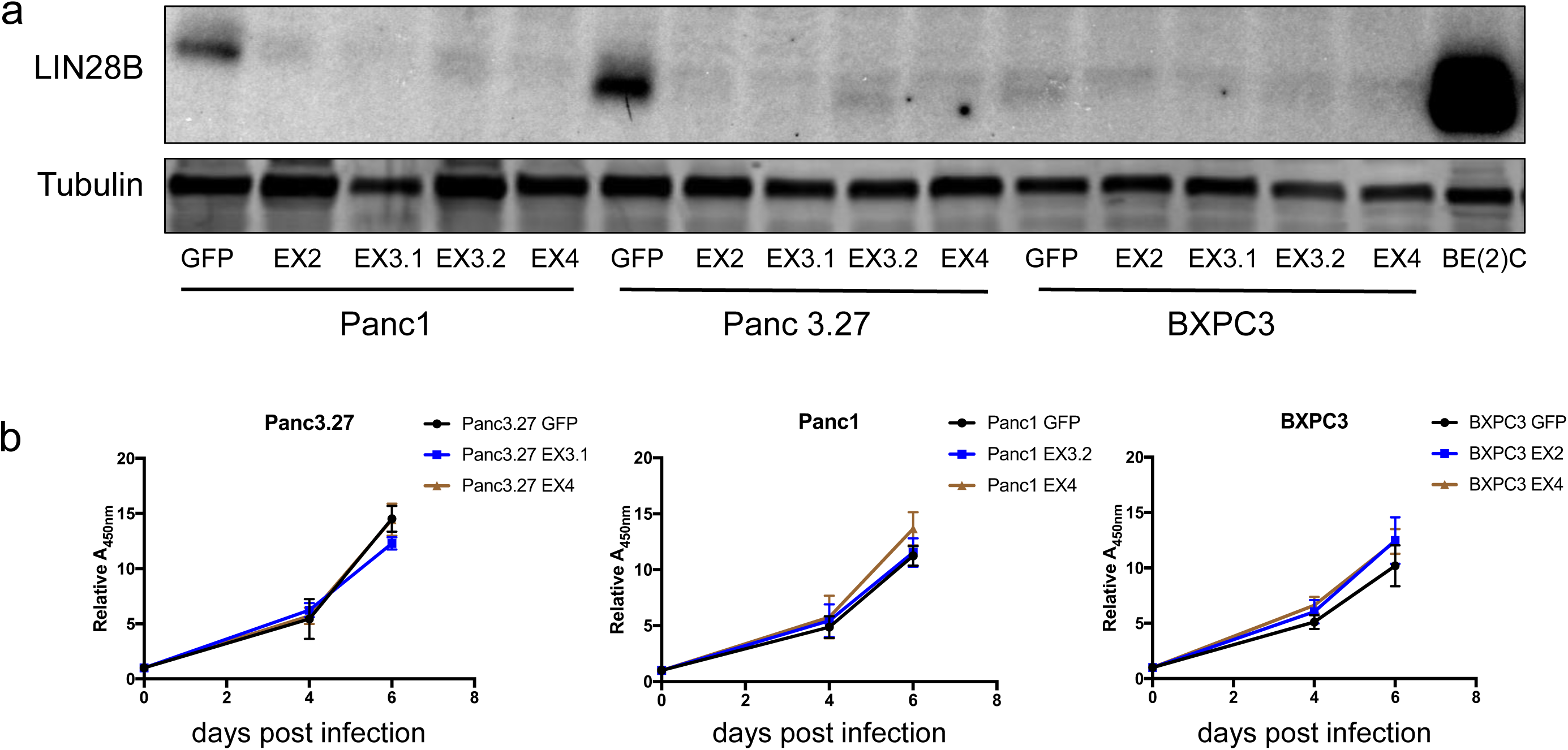
CRISPR/Cas9-mediated genetic loss of *LIN28B* in pancreatic cancer cells. **(a)** Immunoblot for LIN28B in pancreatic cancer cell lines infected with indicated Cas9-gRNA lentivirus. (sgGFP, *GFP*-targeting gRNA control; EX2/3.1/3.2/4, *LIN28B* targeting gRNAs). **(b)** qPCR analysis of relative *let-7* expression in cells from *a*. Fold change relative to *GFP* gRNA controls. **(c)** Cell growth analysis of cells infected as in *a*.

**Figure S3.**
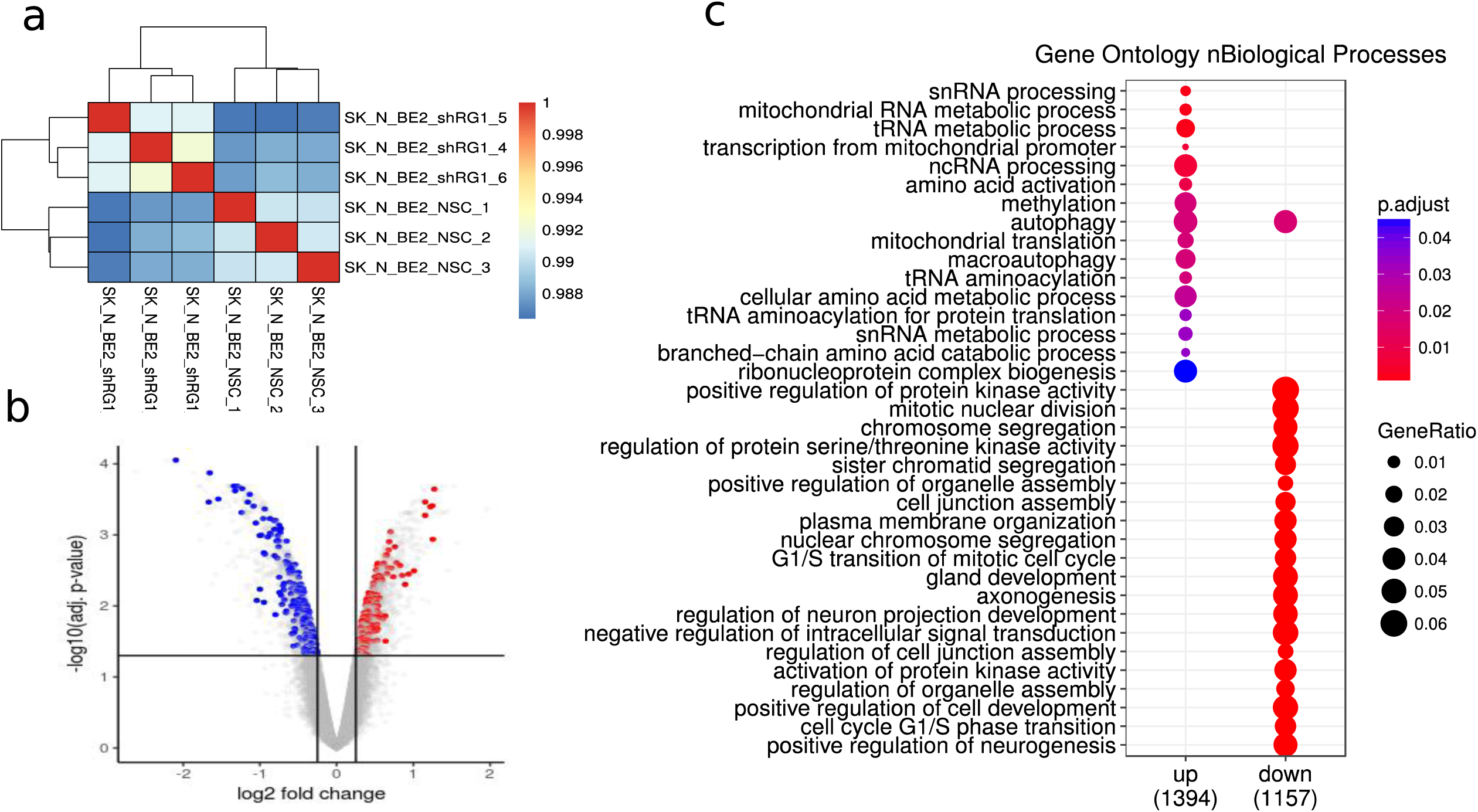
Effect of exogenous shRNA expression on gene expression patterning **(a)** Clustering of Pearson correlation coefficients between gene expression profiles. **(b)** Volcano plots colored by differentially expressed cell cycle-related genes. Blue indicates downregulation; Red indicates upregulation. **(c)** Gene Ontology analysis of differentially expressed genes. Dataset analyzed was generated by Jubierre *et. al.*, 2016.

**Figure S4.**
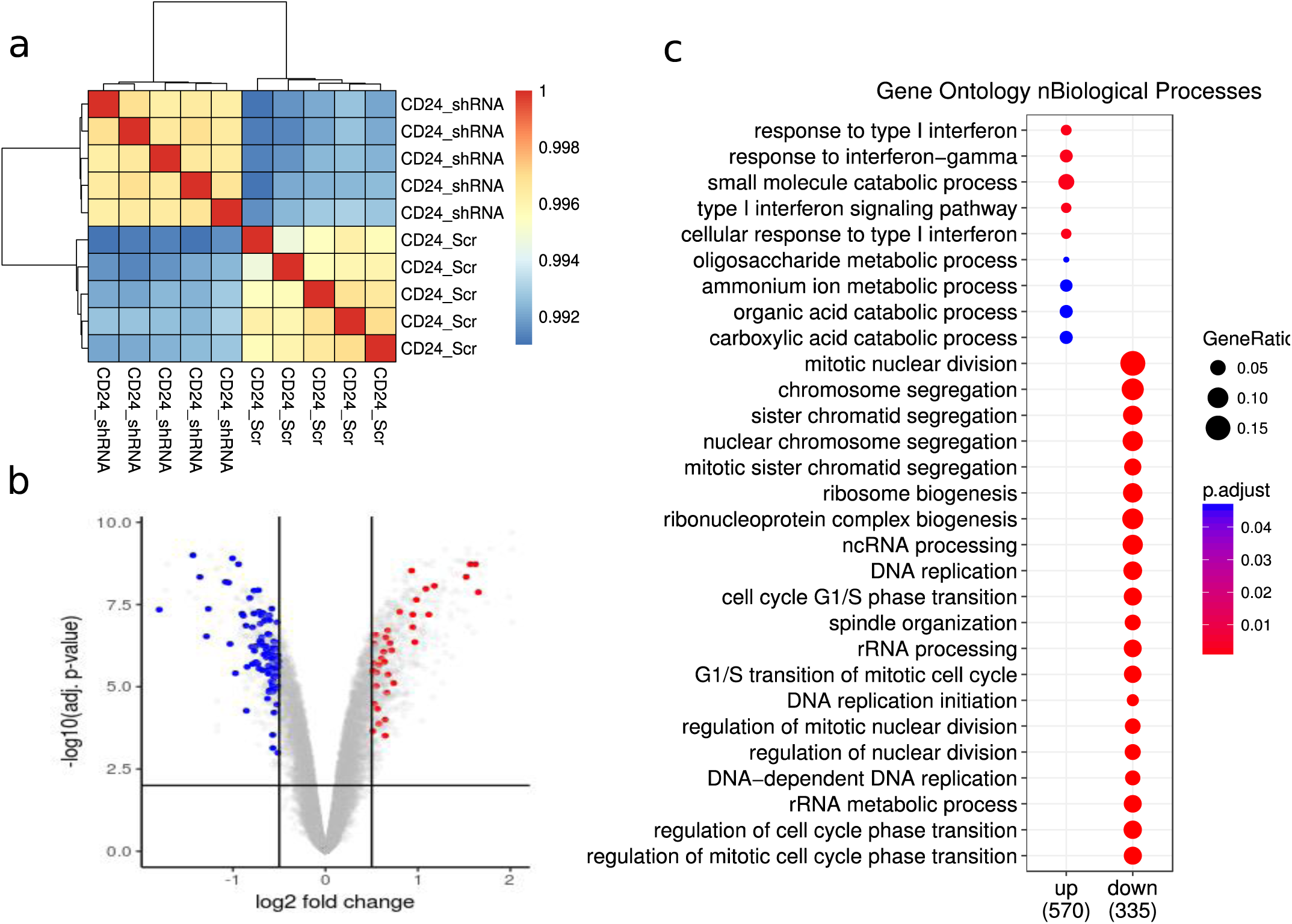
Effect of exogenous shRNA expression on gene expression patterning. **(a)** Clustering of Pearson correlation coefficients between gene expression profiles. **(b)** Volcano plots colored by differentially expressed cell cycle-related genes. Blue, red dots indicate significantly downregulated or upregulated cell cycle related genes, respectively. **(c)** Gene Ontology analysis of differentially expressed genes. Dataset analyzed was generated by Wang *et. al.*, 2017.

**Figure S5.**
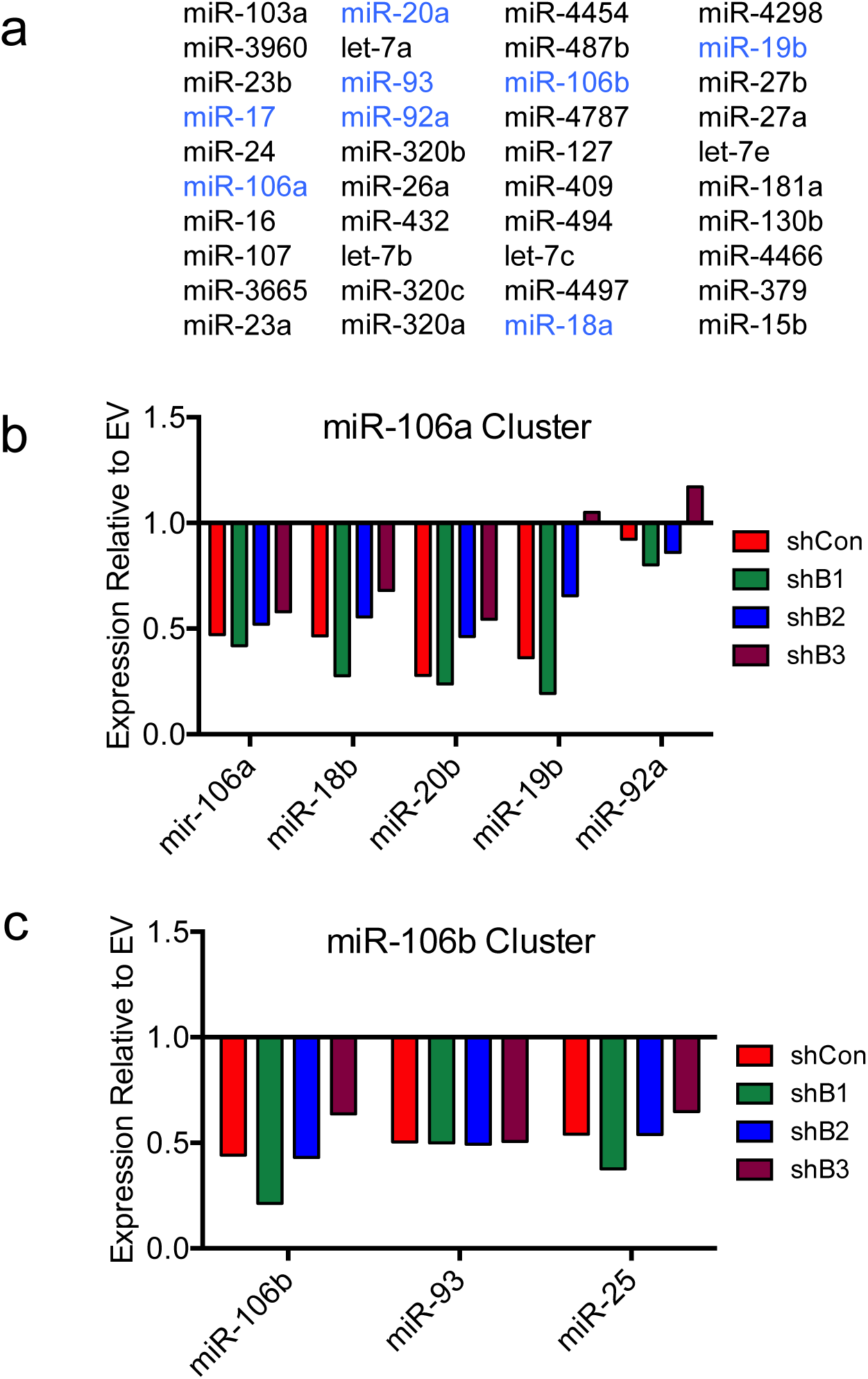
OncomiR microRNA clusters are suppressed by exogenous shRNAs **(a)** List of the 40 most highly expressed microRNAs in BE(2)C EV cells as determined by microRNA microarray. OncomiRs are in blue. **(b,c)** Analysis of relative miR-106a and miR-106b OncomiR cluster expression in cells from infected with the indicated shRNA lentivirus. Values are relative to empty vector control. (EV, empty vector; Con, non-targeting shRNA; B1/B2/B3, *LIN28B*-targeting shRNAs)

**Figure S6.**
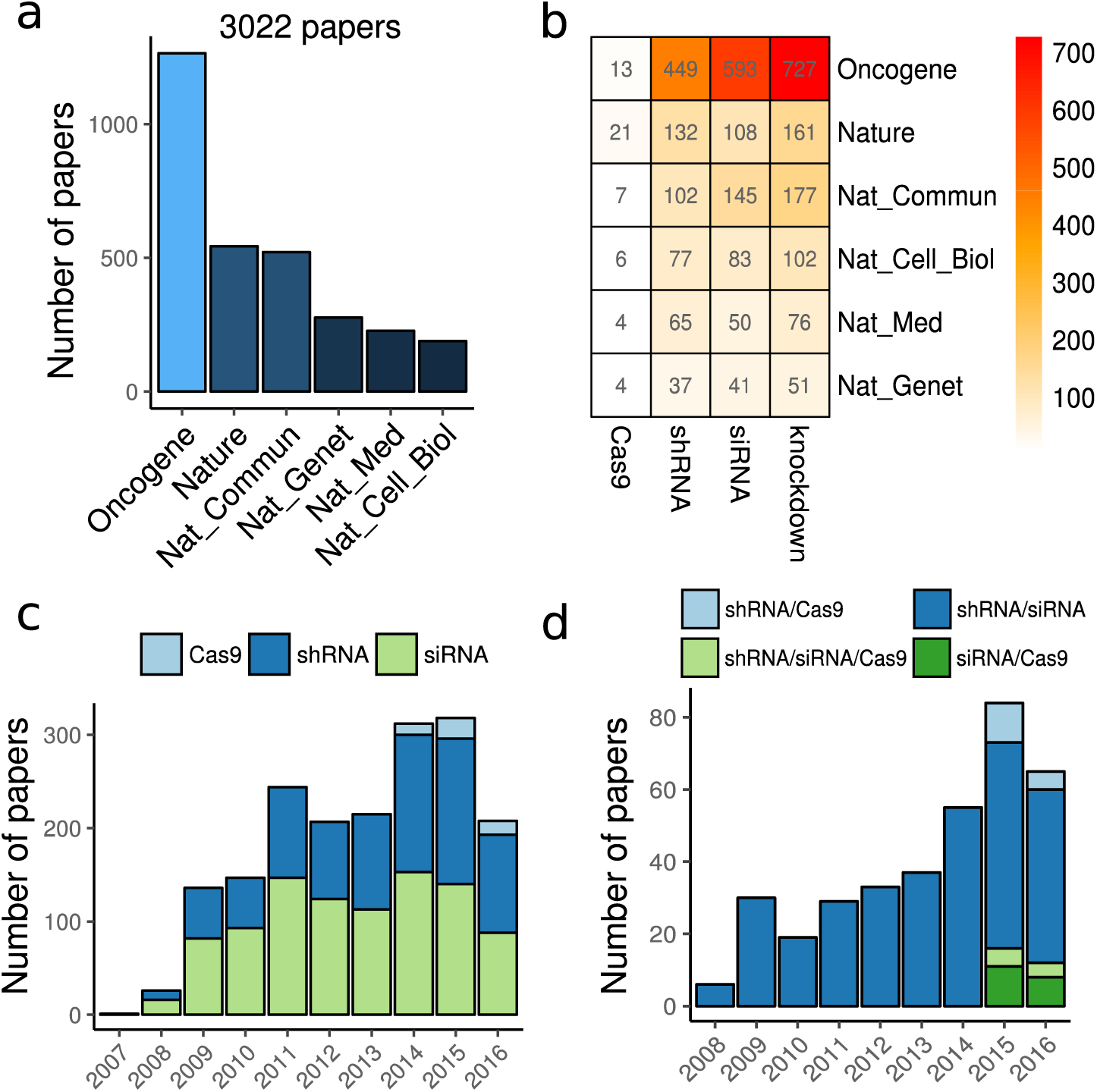
Literature mining of popular knockdown methods. **(a)** Distribution of the number of papers assembled by use of keywords “Cancer” or “Tumor” available in Nature family journal PMC entries. **(b)** Number of papers per journal in *a* containing at least one occurrence of the indicated knockdown methods. **(c)** Number of papers in *b* containing at least one occurrence of indicated knockdown methods from 2007 to 2016. **(d)** Number of papers in *c* containing co-occurrences of selected knockdown methods from 2007 to 2016.

**Figure S7.**
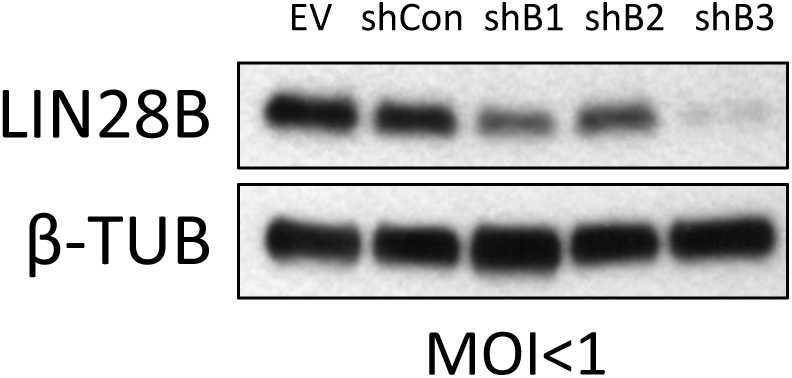
Immunoblot for MYCN and LIN28B in BE(2)C cells infected with the indicated lentiviral constructs (MOI <1) for 7 days.

**Figure S8.**
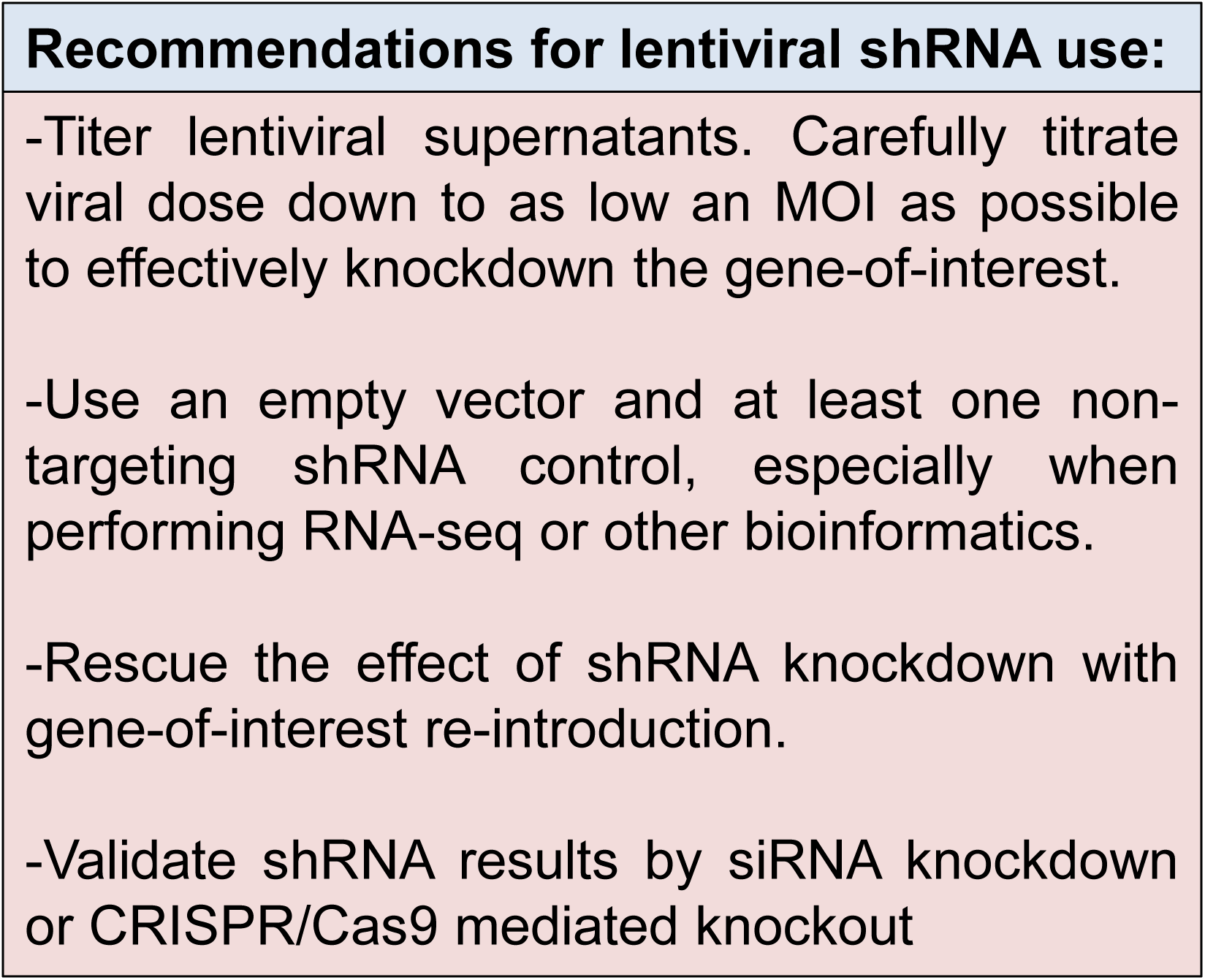
Table of recommendations for shRNA use.

